# Cytotoxic Effects of Multiple Pesticides and their Mixtures on Caco-2 Cells Evaluated by Using MTT and Trypan Blue Assays

**DOI:** 10.64898/2026.09.23.753704

**Authors:** F. Truzzi, E. Tibaldi, S. Dilloo, R. Noferini, D. Sgargi, S. Panzacchi, E. D’Amen, F. Gnudi, G. Dinelli, P.T.J. Scheepers, D. Mandrioli

**Author notes:** Corresponding Authors: Francesca Truzzi, PhD, Junior Assistant Professor, Department of Agricultural and Food Sciences, Alma Mater Studiorum-University of Bologna, Viale Fanin 44, 40127, Bologna, Italy, Daniele Mandrioli, MD, PhD, Former Director, Cesare Maltoni Cancer Research Center, Ramazzini Institute, Via Saliceto, 3, Bentivoglio, Bologna 40010, Italy. co-first authors (F. Truzzi, E. Tibaldi). co-last authors (P.T.J. Scheepers, D. Mandrioli).

## Abstract

**BACKGROUND:** Pesticides are extensively used in agriculture, raising concerns about their potential impact on human health through dietary and environmental exposure. OBJECTIVES: This study evaluated the *in vitro* cytotoxicity of ten commonly used pesticides and their mixtures (lambda-cyhalothrin, cypermethrin, deltamethrin, tebuconazole, glyphosate, acetamiprid, cyprodinil, piperonyl butoxide, fluopyram, and imazalil) on human intestinal Caco-2 cells.

**METHODS:** Cytotoxicity was assessed using the MTT assay, as a measure of metabolic activity, and the trypan blue exclusion test, as an indicator of cell membrane integrity.

**FINDINGS:** Results showed that high concentrations (100 mg L□^1^) of all pesticides significantly reduced cell viability and vitality. Notably, glyphosate and tebuconazole exhibited significant toxicity even at lower concentrations, respectively 0.1 mg L□^1^ and 10 mg L□^1^. Combination treatments (Top 3 and Top 8 pesticide mixtures) retained the cytotoxic effects observed for individual compounds, showing additive (non-synergistic) effects.

**CONCLUSIONS:** Overall, these findings indicate that certain pesticides-based herbicides can exert cytotoxic effects on intestinal cells even at relatively low concentrations and highlight the importance of using the component-based approach in mixture risk assessment for humans. This study was performed as part of the EU SPRINT (Sustainable Plant Protection Transition: A Global Health Approach) project.

## 1. Background

Pesticides are widely used to protect crops from pests, diseases, and weeds, thereby contributing to increased agricultural productivity and food security (Cooper & Dobson, 2007). These chemical agents, while effective in pest control, may lead to unintended human exposure through contaminated food, water, and environmental sources (Damalas & Eleftherohorinos, 2011; EFSA 2023). This widespread use of pesticides in modern agriculture has raised significant concerns regarding their potential toxic effects on human health (Mostafalou & Abdollahi, 2017; Zúñiga-Venegas LA et al 2022). However, despite their benefits, there is growing concern that these compounds may exert adverse effects on human cells, particularly following ingestion (Cimino et al., 2017; Kim et al., 2017).

In this context, assessing cytotoxicity is essential for understanding the potential impact of pesticides on human health. The gastrointestinal tract represents one of the primary sites of exposure, as intestinal epithelial cells are among the first to encounter ingested pesticide residues (Sambuy et al., 2005; Turner JR 2009; EFSA, 2014). Therefore, evaluating the effects of these compounds on intestinal cells is of particular relevance.

The present study focuses on evaluating the *in vitro* toxicity of ten commonly used pesticides lambda-cyhalothrin, cypermethrin, deltamethrin, tebuconazole, glyphosate, acetamiprid, cyprodinil, fluopyram, imazalil, and the synergist piperonyl butoxide. These compounds represent different chemical classes, including pyrethroids, pyrimidines, pyridinyl-ethyl benzamides, azoles, and neonicotinoids, each characterized by distinct mechanisms of action and physicochemical properties (Gilden et al., 2010). The present study is part of the project Sustainable Plant Protection Transition (SPRINT) with support from the EU Horizon 2020 program. A component-based tiered approach was used to assess the effects of individual pesticides and mixtures. The selection of pesticides for the present *in vitro* study, assessed in the first step (Tier 1), was based on the available data from the SPRINT Case Study Sites on over 200 pesticides and in silico modelling, selecting the top 20 pesticides of concern, prioritizing a list of 10 for *in vitro* testing based on occurrence and weighted hazard quotient (wHQ) (see Supplementary material) This selection took into account their widespread use and the potential for their toxic effects to be underestimated in previous studies (Cimino et al., 2017).

Among the available *in vitro* models, Caco-2 cells, derived from human colon adenocarcinoma, are widely used in toxicological studies because they closely mimic the intestinal epithelial barrier, a primary site of chemical absorption and interaction (Sambuy et al., 2005). Several methods are available to assess cell viability and cytotoxicity. The MTT assay measures cellular metabolic activity by quantifying the reduction of tetrazolium salts into insoluble formazan crystals, representing viable cells (Mosmann, 1983). In contrast, the trypan blue exclusion test evaluates cell membrane integrity, distinguishing viable cells from non-viable ones based on dye exclusion (Strober, 2015).

The use of these complementary methods on human intestinal cells, and a more realistic exposure scenario, can provide important insights into the toxicological profiles of pesticides, contributing to a better understanding of their safety and supporting the development of more effective regulatory strategies (Hernández et al., 2013; EFSA, 2019). Therefore, the aim of this study was to investigate the *in vitro* cytotoxicity of ten pesticides on Caco-2 cells, both individually and in combination (Top 3 and Top 8), using MTT and trypan blue assays. Additionally, comparing cytotoxicity data with established ADI values may help identify potential gaps in current safety assessments and support the need for more comprehensive risk evaluation.

## 2. Methods

MTT assay was performed according to ISO 10993-5 (ISO 10993-5 2009).

### 2.1 Materials

The human epithelial colorectal adenocarcinoma Caco-2 cell line (ATCC® HTB-37™) was purchased from a distributor in Italy (Sesto San Giovanni, Milan, Italy). Dulbecco’s Modified Eagle Medium (DMEM), fetal bovine serum (FBS), and penicillin–streptomycin (Pen/Strep) were obtained from Gibco (Waltham, MA, USA).

Imazalil (1-[2-(Allyloxy)-2-(2,4-dichlorophenyl) ethyl] imidazole) (>99% pure), fluopyram (N-[2-[3-Chloro-5-(trifluoromethyl)-2-pyridinyl]ethyl]-2-(trifluoromethyl) benzamide) (>98% pure), piperonyl butoxide (2-(2-Butoxyethoxy) ethyl (6-propylpiperonyl) ether) (>98% pure), cyprodinil (4-Cyclopropyl-6-methyl-N-phenylpyrimidin-2-amine) (> 99% pure), Acetamiprid (N-(6-Chloro-3-pyridylmethyl)-N-cyano-N-methylacetamidine) (>99% pure), glyphosate (N-(Phosphonomethyl) glycine) (>98% pure), tebuconazole (N-(1-(4-Chlorophenyl)-4,4-dimethyl-3-(1H-1,2,4-triazol-1-ylmethyl)-3-pentanol) (>99% pure), deltamethrin (S)-Cyano(3-phenoxyphenyl) methyl (1R,3R)-3-(2,2-dibromoethen-1-yl)-2,2-dimethylcyclopropane-1-carboxylate) (>99% pure), cypermethrin ([Cyano-(3-phenoxyphenyl)methyl]3-(2,2-dichloroethenyl)-2,2-dimethylcyclopropane-1-carboxylate) (>99% pure) and lambda-cyhalothrin (RS)-alpha-cyano-3-phenoxybenzyl 3-(2-chloro-3,3,3-trifluoropropenyl)-2,2,-dimethylcyclopropanecarboxylate) (> 99% pure) were purchased from Merck Life Science S.r.l. (Milan, Italy). All remaining reagents used in the experiments were of analytical grade.

### 2.2. Cell line and culture conditions

Caco-2 cells were cultured in DMEM with 10% FBS and 1% Pen/Strep at 37 °C in a humidified incubator with 5% CO_2_ in tissue culture flasks (75 cm^2^; BD Biosciences, Franklin Lakes, N J, USA), and the culture medium changed every two days. Prior to experimentation, the cells were trypsinized and cell density was evaluated microscopically using a Bürker counting chamber.

### 2.3. Preparation of individual and combined pesticide treatments

Ten pesticides were tested as mono-constituents: acetamiprid, cypermethrin, cyprodinil, deltamethrin, fluopyram, glyphosate, imazalil, lambda-cyhalothrin, piperonyl butoxide, and tebuconazole.

Two mixture treatments were also prepared. The “Top 3” mixture consisted of acetamiprid, glyphosate, and tebuconazole, while the “Top 8” mixture included acetamiprid, cypermethrin, cyprodinil, deltamethrin, glyphosate, lambda-cyhalothrin, piperonyl butoxide, and tebuconazole.

For mono-constituent treatments, each pesticide (5 mg) was dry-mixed with DMEM powder (500 mg) using a mortar and pestle. Stock solutions of 100 mg L□^1^ and 10 mg L□^a^ were prepared by ilution in distilled water. The 100 mg L□^1^ solution was sonicated for 5 min. All stock solutions were filtered through 0.45 μm polyethersulfone (PES) filters before use.

For the Top 3 mixture, each pesticide (10 mg) was dry mixed with DMEM (1000 mg), diluted in distilled water to obtain a 100 mg L□^1^ stock solution, sonicated for 5 min, and filtered (0.45 μm PES). Additional 30 mg L□^1^ stock solutions were prepared by mixing 3 mg of each pesticide with 1 g DMEM powder, followed by dilution, sonication, and filtration. These solutions were combined to obtain final concentrations of 10 mg L□^1^ for each pesticide. Sodium carbonate (Na□CO□) was added to all solutions.

A similar procedure was used for the Top 8 mixture. Each pesticide (10 mg) was dry-mixed with DMEM (1000 mg), diluted to 100 mg L□^1^, sonicated, and filtered. Additional 80 mg L□^1^ stock solutions were prepared and combined to obtain final concentrations of 10 mg L□^1^ for each pesticide.

All stock solutions were subsequently diluted in DMEM to obtain final concentrations of 0.01, 0.1, 1, 10, nd 100 mg L□^1^.

### 2.4 MTT assay

Caco-2 cells (1 × 10□ cells/well) were seeded in 96-well plates (100 μL/well) and incubated for 24 h at 37 °C and 5% CO□. After incubation, the culture medium was replaced with treatment solutions at concentrations of 0.01, 0.1, 1, 10, and 100 mg L□^1^. Untreated cells (DMEM only) were used as the negative control (CTRL), while cells treated with 10% ethanol (ETOH) served as the positive control. After 24 h, cell viability was assessed using the MTT assay. Cells were incubated with MTT solution (1 mg/mL) for 2 h at 37 °C. The solution was then removed by carefully aspirating using a pipette, and formazan crystals were solubilized with isopropanol. Absorbance was measured at 540 nm using a microplate reader (Labsystems Multiskan MS, Thermo Fisher Scientific, Waltham, MA, USA). Results were expressed as percentage viability relative to the untreated control.

### 2.5. Trypan blue exclusion assay

Caco-2 cells (5 × 10□ cells/well) were seeded in 24-well plates and incubated for 24 h. The medium was then replaced with pesticide treatments at concentrations of 0.01, 0.1, 1, 10, and 100 mg L□^1^. Negative and positive control were defined as described for the MTT assay. After 24 h, cells were trypsinized and stained with 0.4% trypan blue (1:1 dilution with cell suspension). Viable cells were counted using a Countess® II FL automated cell counter (Thermo Fisher Scientific, Waltham, MA, USA). Results were expressed as percentage viability relative to the untreated control.

### 2.6 Statistical analysis

All experiments were performed in three independent biological replicates. Within each experiment, six technical replicates per condition were analyzed for the MTT assay, whereas three technical replicates per condition were used for the trypan blue assay. Data are presented as mean ± standard deviation (SD). Statistical analysis was performed using analysis of variance (ANOVA). Based on the ANOVA results, post hoc multiple comparisons were conducted using either Dunn’s test or Dunnett’s test for comparisons with the control group (CTRL), as appropriate.

Statistical significance was assigned at p-value < 0.05. Levels of significance were indicated as follows: * p < 0.05, ** p < 0.01, *** p < 0.001, and **** p < 0.0001 All statistical analyses were performed using Stata version 18 (StataCorp, 2023. *Stata Statistical Software: Release 18*. College Station, TX: StataCorp LLC).

## 3. Results

### 3.1 MTT: Effects of Pesticides, Alone and in Combination, on Caco2 Cells Proliferation

Cell viability of Caco-2 cells was assessed using the MTT assay for each of the 10 individual pesticides as well as for the Top 3 and Top 8 combinations at concentrations of 0.01, 0.1, 1, 10 and 100 mg L□^1^. Results were expressed as percentage viability relative to the untreated control (CTRL) (Table 1). All treatments were performed alongside a positive control consisting of 10% ethanol (ETOH), which reduced cell viability to approximately 40–60% of the untreated CTRL (Figures 1–12A).

**Table 1.** Summary table comparing MTT cell viability values (%) in Caco-2 cells exposed to individual pesticides and the Top 3 and Top 8 mixtures at concentrations ranging from 0.01 to 100 mg L□^1^. Cell viability is expressed as a percentage relative to the untreated control (CTRL).ss

| Pesticide treatment | 0.01 | 0.1 | 1 | 10 | 100 |
| --- | --- | --- | --- | --- | --- |
| cyhalothrin | 97,232 | 114,527 | 120,718 | 110,174 | **42,741** |
| cypermethrin | 109,506 | 110,471 | 87,039 | 100,319 | **43,131** |
| deltamethrin | 108,901 | 116,205 | 128,173 | 125,730 | **39,782** |
| **tebuconazole** | **99,951** | **88,854** | **87,904** | **66,398** | **31,641** |
| glyphosate | 100,222 | 80,003 | 88,102 | 95,093 | **22,219** |
| acetamiprid | 124,664 | 146,926 | 100,907 | 104,369 | **35,102** |
| cyprodinil | 162,857 | 97,486 | 94,898 | 101,266 | **32,393** |
| piperonyl butoxide | 113,800 | 112,924 | 121,822 | 86,582 | **24,333** |
| fluopyram | 114,948 | 113,584 | 99,358 | 99,214 | **45,990** |
| imazalil | 144,094 | 95,230 | 109,702 | 96,755 | **27,196** |
| Top 3 | 127,963 | 132,242 | 147,185 | 92,730 | **41,600** |
| Top 8 | 124,091 | 95,470 | 118,082 | 101,096 | **13,244** |

**FIGURE 1.**
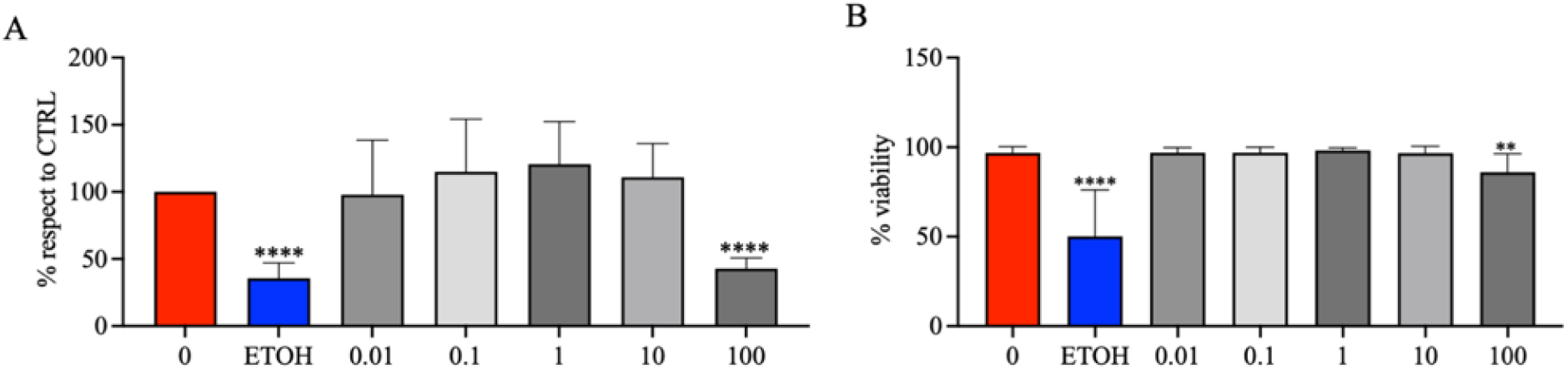
MTT (proliferation) assay and Trypan Blue (viability) assay: cyhalothrin. Effects of pure lambda cyhalothrin, at the dose range 0.01, 0.1, 1.0, 10 and 100 mg L^−1^ on cell proliferation (A) and on cell viability (B) of Caco-2 cells. Data are expressed as mean value (± st. dev.) (% compared to control). Pairwise comparison based on Anova (Dunnett test): *P < 0.05; **P < 0.01; ***P < 0.001; ****P < 0.0001.

**FIGURE 2.**
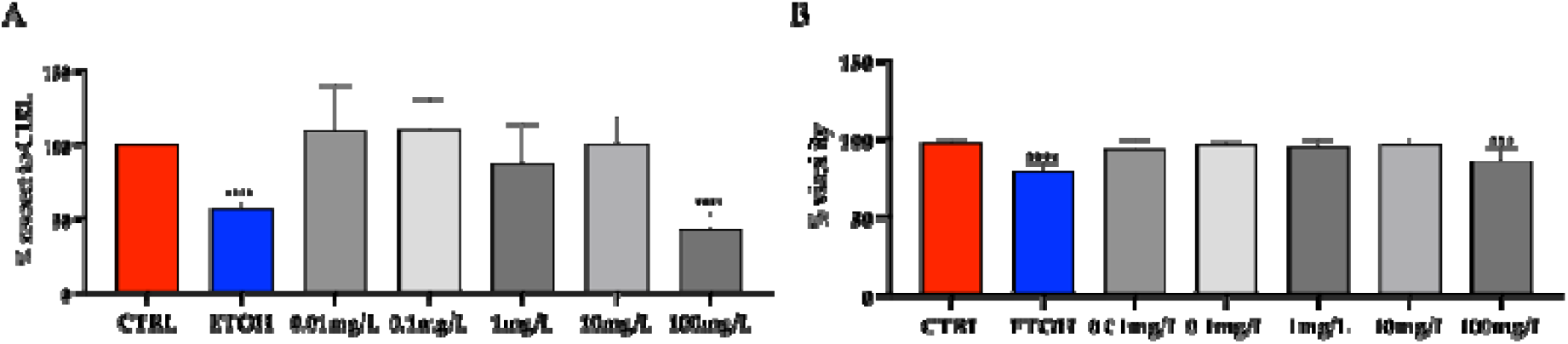
MTT (proliferation) assay and Trypan Blue (viability) assay: cypermethrin. Effects of pure cypermethrin, at the dose range 0.01, 0.1, 1.0, 10 and 100 mg L^−1^ on cell proliferation (A) and on cell viability (B) of Caco-2 cells. Data are expressed as mean value (± st. dev.) (% compared to control). Pairwise comparison based on Anova (Dunnett test): *P < 0.05; **P< 0.01; ***P < 0.001; ****P < 0.0001.

**FIGURE 3.**
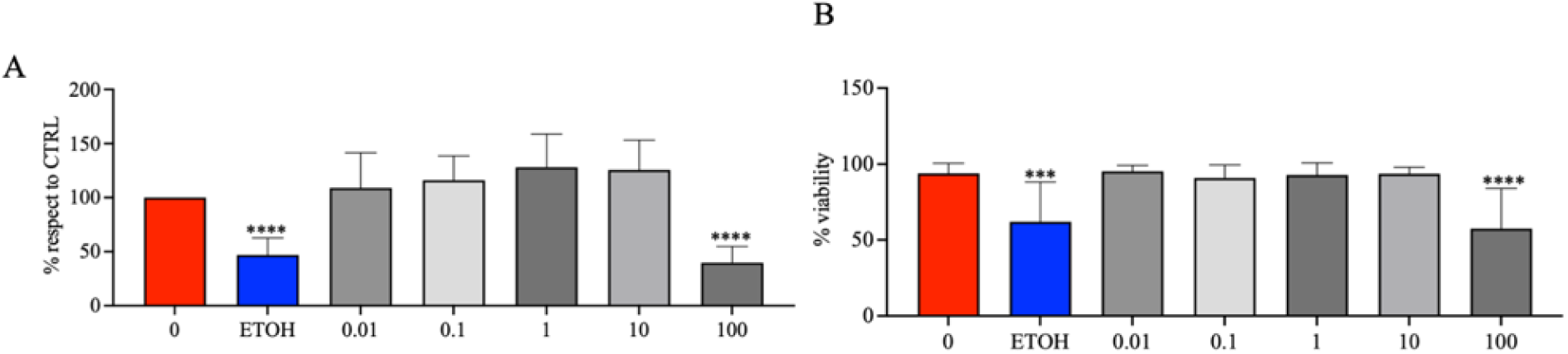
MTT (proliferation) assay and Trypan Blue (viability) assay: deltamethrin. Effects of pure deltamethrin, at the dose 0.01, 0.1, 1.0, 10 and 100 mg L^−1^ on cell proliferation (A) and on cell viability (B) of Caco-2 cells. Data are expressed as mean value (± st. dev.) (% compared to control). Pairwise comparison based on Anova (Dunnett test): *P < 0.05; **P < 0.01; ***P < 0.001; ****P < 0.0001.

**FIGURE 4.**
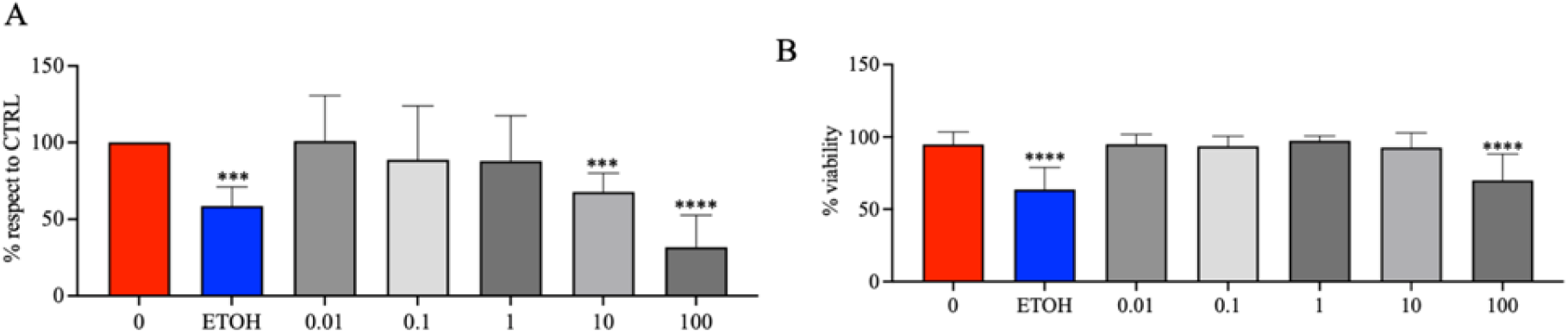
MTT (proliferation) assay and Trypan Blue (viability) assay: tebuconazole. Effects of pure tebuconazole, at the dose 0.01, 0.1, 1.0, 10 and 100 mg L^−1^ on cell proliferation (A) and on cell viability (B) of Caco-2 cells. Data are expressed as mean value (± st. dev.) (% compared to control). Pairwise comparison based on Anova (Dunnett test): *P < 0.05; **P < 0.01; ***P < 0.001; ****P < 0.0001.

A statistically significant reduction in cell viability was observed at the highest concentration tested (100 mg L□^1^) for all individual test compounds (Figures 1–10A) as well as for both combination treatments, Top 3 (Figure 11A) and Top 8 (Figure 12A) (p < 0.0001). This concentration was the only one consistently associated with a marked decrease in cell viability across all tested conditions. At 100 mg L□^1^, cell viability did not exceed 20% of the untreated CTRL for glyphosate (Figure 5A), piperonyl butoxide (Figure 8A), imazalil (Figure 10A), and the Top 8 mixture (Figure 12A). Viability remained below 25% of the untreated CTRL for tebuconazole (Figure 4A) and cyprodinil (Figure 7A). For the remaining pesticides and for the Top 3 mixture (Figure 11A), cell viability values at 100 mg L□^1^ were comparable to those observed for the positive control (10% ETOH), indicating a strong cytotoxic effect. At lower concentrations, limited effects on cell viability were observed. Notably, glyphosate significantly reduced cell viability at 0.1 mg L□^1^ (p < 0.01; Figure 5A), while tebuconazole induced a significant reduction in cell viability at 10 mg L□^1^ (p < 0.001; Figure 4A). For all other pesticides and for both combination treatments, significant reductions in cell viability were observed at concentrations equal to 100 mg L□^1^ (Table 1).

**FIGURE 5.**
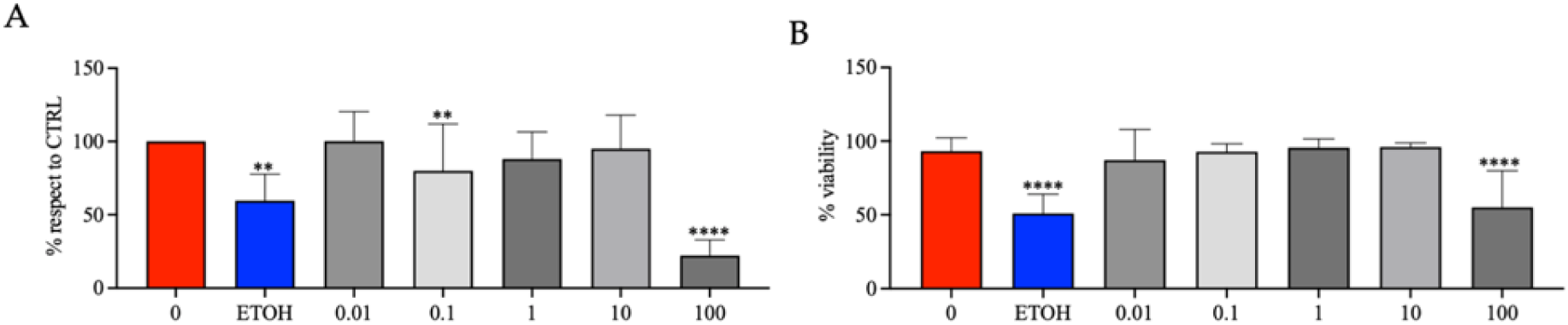
MTT (proliferation) assay and Trypan Blue (viability) assay: glyphosate. Effects of pure glyphosate, at the dose range 0.01, 0.1, 1.0, 10 and 100 mg L^−1^ on cell proliferation (A) and on cell viability (B) of Caco-2 cells. Data are expressed as mean value (± st. dev.) (% compared to control). Pairwise comparison based on Anova (Dunnett test): *P < 0.05; **P < 0.01; ***P < 0.001; ****P < 0.0001.

**FIGURE 6.**
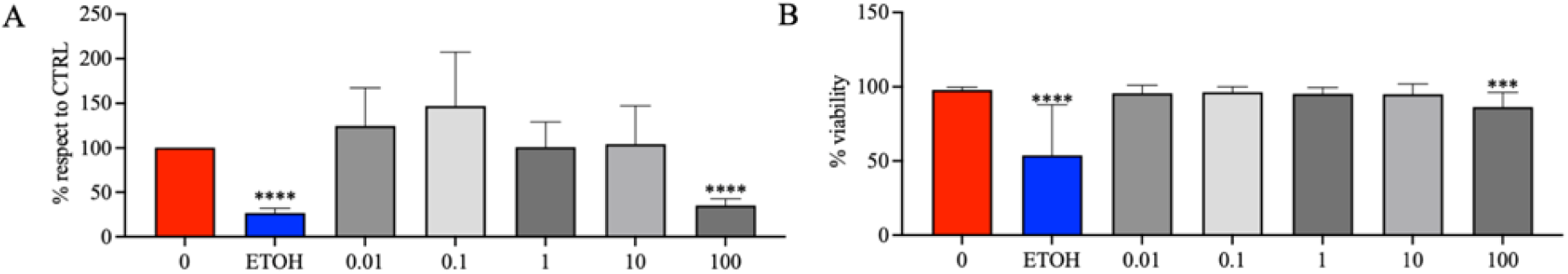
MTT (proliferation) assay and Trypan Blue (viability) assay: acetamiprid. Effects of pure acetamiprid, at the dose range 0.01, 0.1, 1.0, 10 and 100 mg L^−1^ on cell proliferation (A) and on cell viability (B) of Caco-2 cells. Data are expressed as mean value (± st. dev.) (% compared to control). Pairwise comparison based on Anova (Dunnett test): *P < 0.05; **P < 0.01; ***P < 0.001; ****P < 0.0001.

**FIGURE 7.**
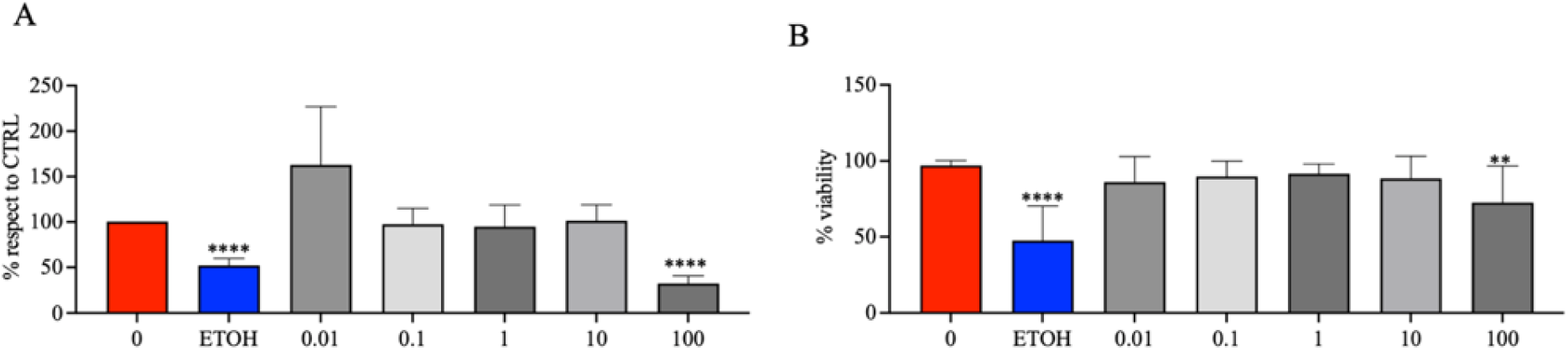
MTT (proliferation) assay and Trypan Blue (viability) assay: cyprodinil. Effects of pure cyprodinil, at the dose range 0.01, 0.1, 1.0, 10 and 100 mg L^−1^ on cell proliferation (A) and on cell viability (B) of Caco-2 cells. Data are expressed as mean value (± st. dev.) (% compared to control). Pairwise comparison based on Anova (Dunnett test): *P < 0.05; **P < 0.01; ***P < 0.001;****P < 0.0001.

**FIGURE 8.**
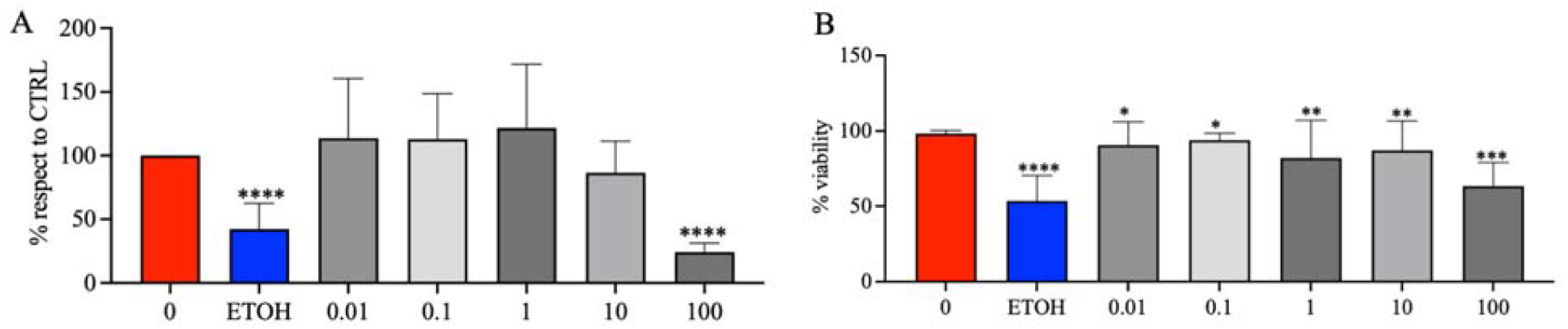
MTT (proliferation) assay and Trypan Blue (viability) assay: piperonyl butoxide. Effects of pure piperonyl butoxide, at the dose range 0.01, 0.1, 1.0, 10 and 100 mg L^−1^ on cell proliferation (A) and on cell viability (B) of Caco-2 cells. Data are expressed as mean value (± st. dev.) (% compared to control). Pairwise comparison based on Anova (Dunnett test): *P < 0.05; **P< 0.01; ***P < 0.001; ****P < 0.0001.

**FIGURE 9.**
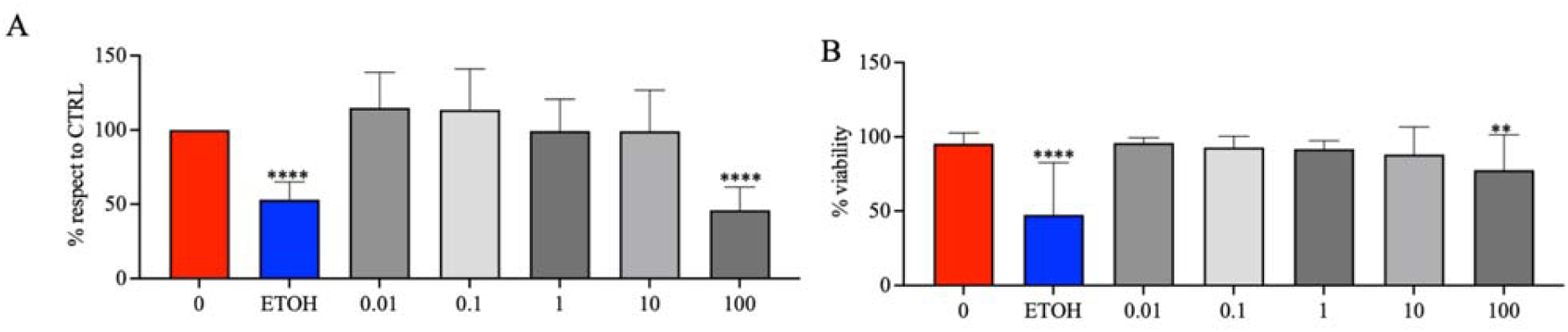
MTT (proliferation) assay and Trypan Blue (viability) assay: fluopyram. Effects of pure fluopyram, at the dose range 0.01, 0.1, 1.0, 10 and 100 mg L^−1^ on cell proliferation (A) and on cell viability (B) of Caco-2 cells. Data are expressed as mean value (± st. dev.) (% compared to control). Pairwise comparison based on Anova (Dunnett test): *P < 0.05; **P < 0.01; ***P < 0.001; ****P < 0.0001.

**FIGURE 10.**
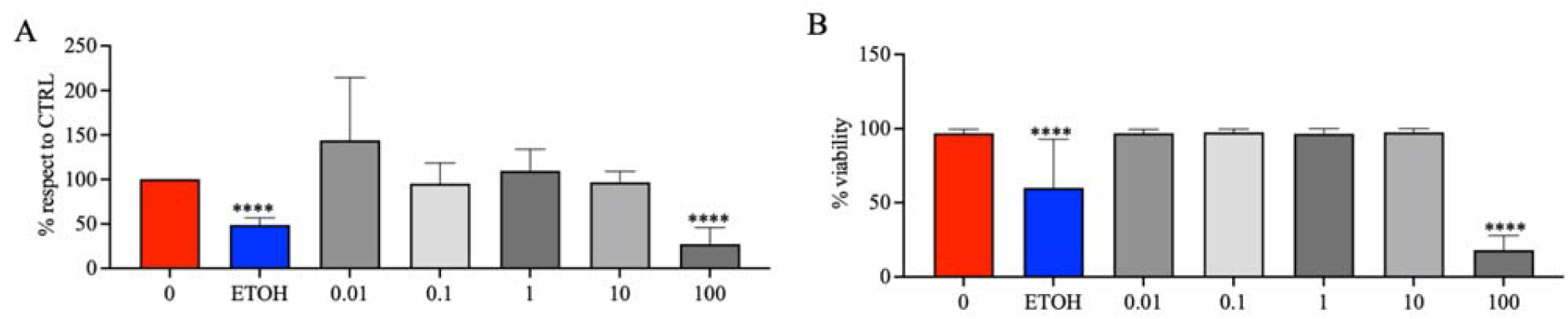
MTT (proliferation) assay and Trypan Blue (viability) assay: imazalil. Effects of pure imazalil, at the dose range 0.01, 0.1, 1.0, 10 and 100 mg L^−1^ on cell proliferation (A) and on cell viability (B) of Caco-2 cells. Data are expressed as mean value (± st. dev.) (% compared to control). Pairwise comparison based on Anova (Dunnett test): *P < 0.05; **P < 0.01; ***P < 0.001 ; ****P < 0.0001.

**FIGURE 11.**
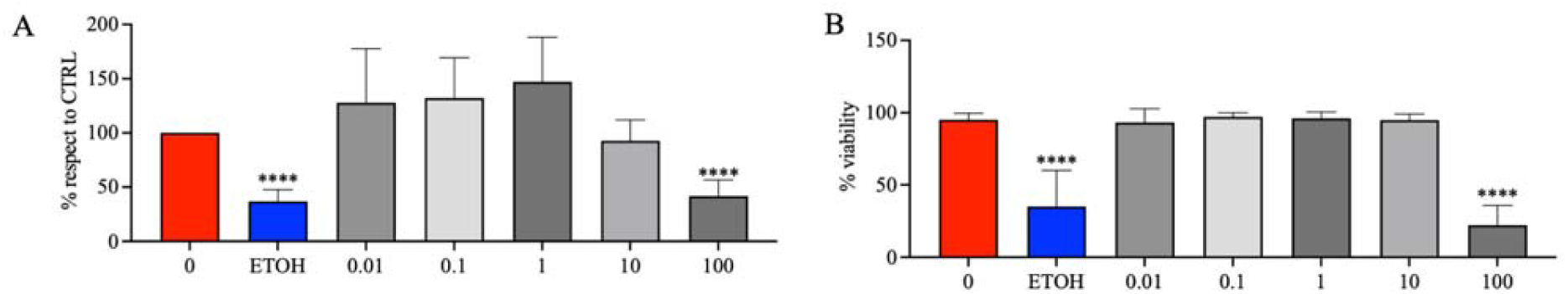
MTT (proliferation) assay and Trypan Blue (viability) assay: Top 3. Effects of Top 3 combination of pesticide (pure acetamiprid, tebuconazole, glyphosate), at the dose range 0.01, 0.1, 1.0, 10 and 100 mg L^−1^ on cell proliferation (A) and on cell viability (B) of Caco-2 cells. Data are expressed as mean value (± st. dev.) (% compared to control). Pairwise comparison based on Anova (Dunnett test): *P < 0.05; **P < 0.01; ***P < 0.001; ****P < 0.0001.

**FIGURE 12.**
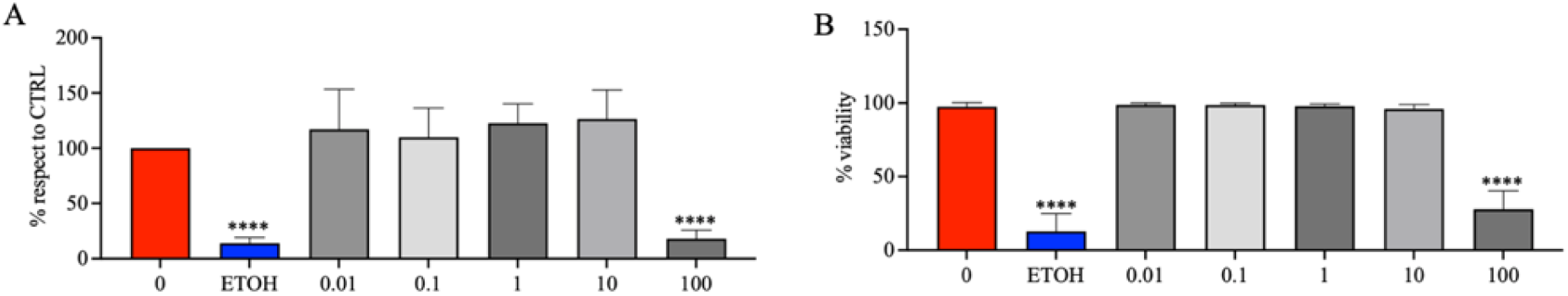
MTT (proliferation) assay and Trypan Blue (viability) assay: Top 8. Effects of Top 8 combination of pesticide (pure lambda cyhalothrin, cypermethrin, deltamethrin, tebuconazole, glyphosate, acetamiprid, cyprodinil, piperonyl butoxide), at the dose range 0.01, 0.1, 1.0, 10 and 100 mg L^−1^ on cell proliferation (A) and on cell viability (B) of Caco-2 cells. Data are expressed as mean value (± st. dev.) (% compared to control). Pairwise comparison based on Anova (Dunnett test): *P < 0.05; **P < 0.01; ***P < 0.001; ****P < 0.0001.

The individual pesticides and mixtures maintain cell viability close to or above control levels up to 10 mg L□^1^, sometimes inducing proliferative responses. Based on the data provided, tebuconazole is the most toxic individual compound at lower to intermediate concentrations, while the Top 8 mixture induces a significantly higher cytotoxic effect than the Top 3 mixture at the highest concentration. Tebuconazole shows the lowest cell viability at 10 mg L□^1^ (66.40%) and steadily decreases from 0.1 mg L□^1^ (88.85%). Glyphosate exhibits the lowest cell viability at 0.1 mg L□^1^ (80.00%), though it slightly recovers at 10 mg L□^1^. At highest dose (100 mg L□^1^) Top 8 shows severe synergistic toxicity, collapsing viability to 13.24%, compared to 41.60% for Top 3 (Table 1). Overall, these results indicate that the cytotoxic effects of the tested pesticides on Caco-2 cells are primarily concentration-dependent, with significant reductions in cell viability occurring predominantly at the highest concentration tested, moreover only selected compounds, such as glyphosate and tebuconazole, exhibited effects at lower concentrations.

### Trypan Blue: Effects of Pesticides, Alone and in combination, on Caco2 Cells Viability

Cell viability of Caco-2 cells was evaluated using the trypan blue exclusion assay for each of the 10 individual pesticides as well as for the Top 3 and Top 8 combinations at concentrations of 0.01, 0.1, 1, 10, and 100 mg L□^1^. Results were expressed as percentage viability relative to the untreated control (CTRL). The positive control (10% ethanol, ETOH) reduced cell vitality to approximately 50–60% of the untreated CTRL (Figures 1–12B). A statistically significant reduction in cell vitality was observed at 100 mg L□^1^ for several pesticides, including deltamethrin (Figure 3B), tebuconazole (Figure 4B), glyphosate (Figure 5B), imazalil (Figure 10B), and for both combination treatments, Top 3 (Figure 11B) and Top 8 (Figure 12B) (p < 0.0001). Significant reductions were also observed at the same concentration for cypermethrin (Figure 2B), acetamiprid (Figure 6B), and piperonyl butoxide (Figure 8B) (p < 0.001), as well as for lambda-cyhalothrin (Figure 1B), cyprodinil (Figure 7B), and fluopyram (Figure 9B) (p < 0.01). At 10 mg L□^1^, a significant reduction in cell vitality was observed only for piperonyl butoxide (p < 0.01; Figure 8B). For all other pesticides and for both combination treatments, no significant effects were detected at lower concentrations compared to the untreated CTRL. Overall, the trypan blue assay confirmed the dose-dependent effects observed in the MTT assay, with significant reductions in cell vitality occurring predominantly at the highest concentration tested, and limited effects at lower concentrations.

## 4. Discussion

The results of this study demonstrated that the toxic effects of the tested pesticides on Caco-2 cells are predominantly evident at the highest concentration tested (100 mg L□^1^), significantly reducing both cell viability and vitality in the MTT and trypan blue assays, for all individual pesticides and their combinations (Top 3 and Top 8). The concordance between the two assays further strengthens the reliability of the observed cytotoxic effects, as they assess complementary cellular endpoints. Notably, the 100 mg L□^1^ treatment resulted in cell viability decreasing below 20–25% for several pesticides, including glyphosate, imazalil, tebuconazole, and cyprodinil, and the synergist piperonyl butoxide, indicating a strong cytotoxic effect at this concentration. These results are consistent with previous studies highlighting the dose-dependent toxicity of pesticides on human cell lines (Hernández et al., 2013; Truzzi et al. 2021; Silva et al., 2022). Of the tested pesticides, glyphosate and tebuconazole emerged as particularly potent, exhibiting significant toxic effects even at lower concentrations of 0.1 mg L□^1^ and 10 mg L□^1^, respectively. Glyphosate, a widely used herbicide, reduced cell viability significantly also at 0.1 mg L□^1^ (p < 0.01), while tebuconazole, a triazole fungicide, showed a similar effect at 10 mg L□^1^ (p < 0.001). Importantly, these concentrations approach levels that may be relevant for environmental and dietary exposure, raising concerns about potential implications of real-world exposures. Different studies have highlighted that low-dose pesticide exposure may induce relevant effects, including oxidative stress, endocrine disruption, and cancer (Myers et al., 2016; Mesnage & Antoniou, 2017; Vandenberg et al., 2012; Panzacchi et al, 2025). Similarly, tebuconazole’s ability to disrupt cellular membranes or interfere with enzymatic processes may contribute to its observed toxicity (Zubrod et al., 2019). These mechanistic insights support the biological plausibility of the effects observed in this study.

In contrast, pesticides such as lambda-cyhalothrin and fluopyram exhibited a less pronounced impact on Caco-2 cells at lower concentrations but maintained the significance at the highest dose. This suggests that these compounds may be better tolerated by intestinal cells, possibly due to differences in their chemical structures, modes of action, and metabolic pathways (Naravaneni et al., 2005; Ilboudo et al., 2014). However, the absence of effects at lower doses should be interpreted with caution, as sub-lethal cellular alterations not captured by these assays may still occur. Indeed, emerging evidence indicates that low-dose exposures may alter gene expression, barrier integrity, and inflammatory responses without overt cytotoxicity (Zhou M et al, 2021; Kortenkamp A. 2008). However, it is important to note that even these pesticides showed significant cytotoxicity at the highest concentration (100 mg L□^1^), underscoring the importance of dose-dependent effects in toxicological assessments. This highlights the need to consider a wide range of concentrations when evaluating pesticide safety.

The combined treatments to the Top 3 and Top 8 mixtures revealed that the toxic effects observed for individual pesticides were preserved when administered in mixtures, showing an additive (non-synergistic) response. This suggests that the combined exposure to multiple pesticides, even at lower concentrations of single test compounds, could still pose a significant risk to intestinal health.

This finding reflects realistic exposure scenarios, where humans are more likely to be exposed to mixtures rather than single compounds.

In summary, these findings underscore the importance of monitoring pesticide concentrations in biological environments, particularly for substances like glyphosate and tebuconazole, which exhibit significant toxicity even at low concentrations. No synergistic interactions were evident in either the mixture of 3 or 8 prioritized pesticides. This study provides important information on the cytotoxicity of different widely used pesticides and confirms the validity of a component-based approach in genotoxicity assessment of pesticide mixtures. Future studies should focus on elucidating the mechanisms underlying the observed cytotoxicity, investigating the effects of chronic low-dose exposures. Additionally, integrating omics-based approaches and barrier function assays could provide further mechanistic insights into subtle cellular alterations. The limitations are related to the extrapolation of these results to the *in vivo* situation where the integrity of the microenvironment of the complete intestinal tissue will determine the toxicity of the tested compounds including metabolism of pesticides formed by the gut microflora.

## Conclusion

This study demonstrates that commonly used pesticide can exert significant cytotoxic effects on human intestinal Caco-2 cells, with certain compounds such as glyphosate and tebuconazole showing significant effects even at lower doses. The consistency between MTT and trypan blue assays strengthens the robustness of these findings. Importantly, pesticide mixtures (Top 3 and Top 8) preserved the cytotoxic effects observed for individual compounds, highlighting the potential risks associated with combined exposures. These results are particularly relevant in real-world contexts, where humans are simultaneously exposed to multiple pesticide residues through diet and the environment. The observed effects at relatively low concentrations for specific pesticides underscore the need for further investigation into chronic exposure and mixture toxicity using human-relevant models. This study also confirms the validity of a component-based approach in the assessment of pesticide mixtures for toxicity.

## CRediT authorship contribution statement

**FT:** conceptualization, data curation and formal analysis, investigation, visualization, writing the original work, reviewing and editing

**ET:** conceptualization, data curation and formal analysis, investigation, visualization, reviewing and editing

**SD:** investigation

**RN**: investigation

**DS:** data curation and formal analysis, reviewing and editing

**SP:** data curation and formal analysis, reviewing and editing

**EDA:** investigation

**FG:** investigation

**GD:** reviewing and editing

**PTJS:** reviewing and editing, supervision, project administration and resources for study conduct.

**DM** conceptualization, investigation, visualization, writing the original work, reviewing and editing, supervision, project administration and resources for study conduct.

## Data availability

No datasets were generated or analyzed during the current study

## Acknowledgement

We acknowledge the contribution of Hans Mol to the development and analyses of the data of the in silico modelling

