## Supplementary Material 1 for "Cytotoxic Effects of Multiple Pesticides and their Mixtures on Caco-2 Cells Evaluated by Using MTT and Trypan Blue Assays"

### Selection of Pesticides for toxicological studies

Widespread use of Pesticides leads to exposure from dietary sources and from environmental contamination resulting in uptake of pesticide residues by consumers, farmers and families living near agricultural fields (neighbours). A procedure was developed to prioritize Pesticides. for toxicology testing. The priority setting was done using a stepwise approach: occurrence and exposure assessment (A), hazard assessment (B) and in the third step the synthesis (C) of the results of the two previous steps A and B. For assessment of occurrence, we used existing data on food residue analysis and new data from the SPRINT field study on human exposure (pesticide residues in blood, (urine) and feces) and occurrence in the environment (primarily environmental concentration data from farms). Occurrence data were converted to exposure expressed as body burden in mg per kg bodyweight per day (mg/kg/d). For this subset hazard quotients (HQ) were calculated based on median exposure levels form biomonitoring data (blood and feces) and acceptable daily intake (ADI) and acceptable operator exposure limit (AOEL) used as a reference of risk for consumers and farmers and neighbours, respectively. We calculated HQs for these two populations. HQs were weighted based on frequency of occurrence of single pesticide divided by the total number of Pesticides in a give sample matrix (blood, (urine) and feces) to account for Pesticides with very low detection rates. The weigted HQs were then used to rank pesticides for the two sub-populations of interest.

The first step resulted in a reduction from a long list of 206 Pesticides to a short list of 10 pesticides based on existing and new occurrence data. HQs plotted against exposure data in a risk matrix. Next, HQs were weighted by the detection frequencies in the human samples. This resulted in a ranking of 14 pesticide active ingredients for which sufficient data were available. For the remaining substance occurrence and hazard data were insufficient. The short list of 14 pesticide included: lambda-cyhalothrin, cypermethrin, deltamethrin, folpet, glyphosate, acetamiprid, cyprodinil, piperonyl butoxide (synergist), fluopyram, imazalil, pendimethalin, trifloxystrobin, fluonicamid, and fludioxonil. Despite the differences between HQs based on

ADIs (derived from consumer exposures) and those derived from OELs (from farmer's and neighbour's exposures) good agreement was observed in the rank numbers

Ranking of the 20 test compounds and identification of the relevant mixtures was done using the procedure shown in **Figure S1**.

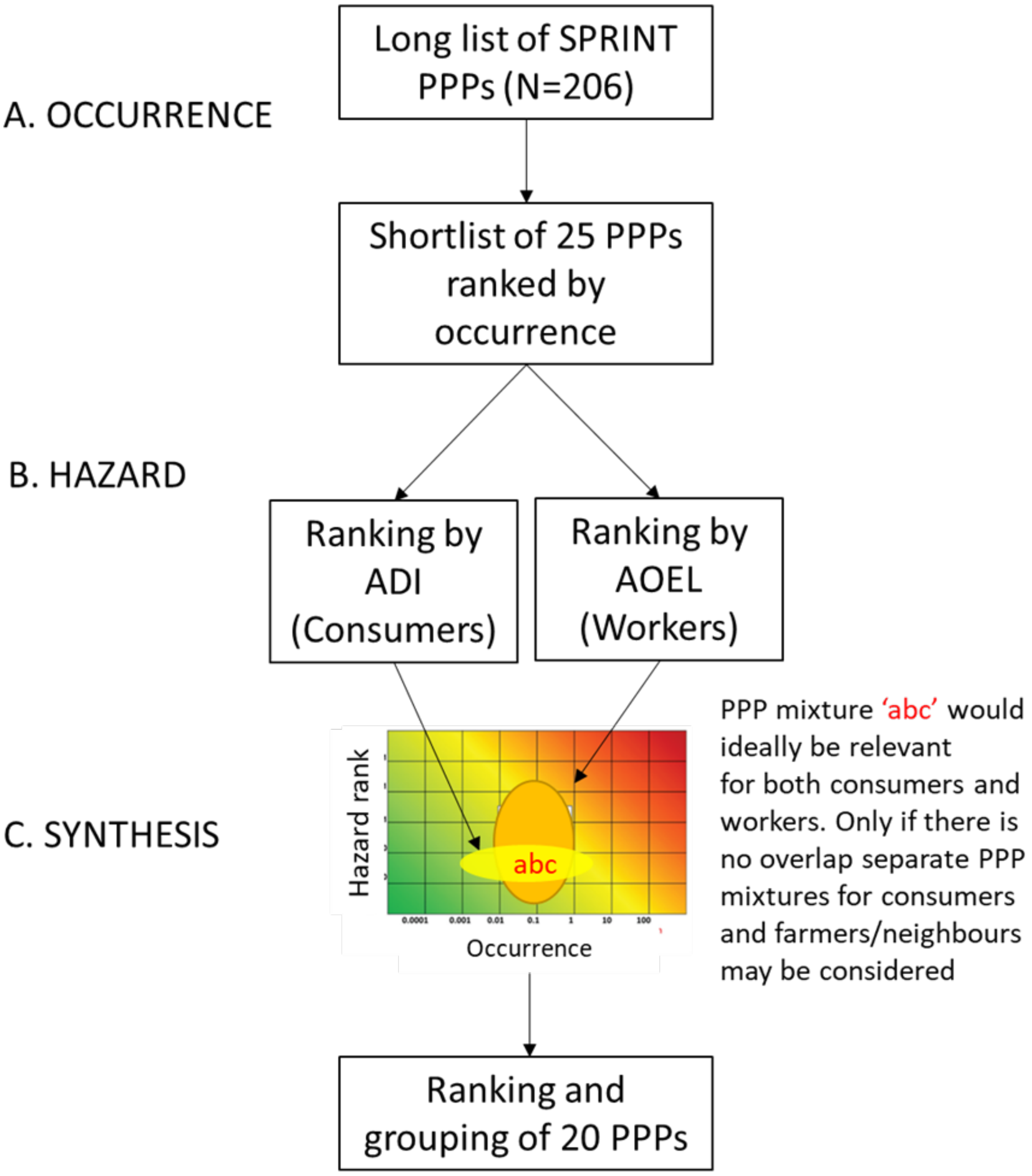


**Figure S1.** Occurrence and exposure (A) is based on approval status and the occurrence in environmental and biological matrices. Hazard (B) is based on the acceptable daily intake (ADI) for the general public and the acceptable operator exposure level (AOEL) relevant to occupational exposure scenarios. In the third step the synthesis (C) is performed by a risk plot.

### Step A: Occurrence and exposure

The selection process for human health considerations started with an assessment of occurrence and exposure. The rationale for this is that without information on exposure, it will not be possible to assess the impact on human health in the context of the SPRINT project. The starting point for occurrence assessment was the SPRINT list of Pesticides established for the field study (Silva et al., 2021). The background on the establishment of this target list has been described in SPRINT report “Pesticide component selection chemical analysis” (Mol, 05.05.2021), and a summary of this has been included in the publication of the SPRINT study protocol (Silva et al., 2021).

For human exposure, both internal and external exposure were considered. Internal exposure data was obtained through human biomonitoring (HBM), here data of pesticide (biomarkers of exposure) in blood, urine and feces. External exposure data included food (oral intake), air and (fine) indoor dust (inhalation route, possibly dermal), and occurrence in other environmental matrices (potentially oral/dermal exposure). An indication for (potential) exposure was based on the overall data generated in SPRINT, rather than at an individual CSS level, because the number of samples per CSS was rather low in several cases. Data from outside SPRINT were considered in case data from SPRINT CSSs were not (yet) available by July 2022.

In Table S1 an overview is given for all data sources included for potential human exposure, with the number of samples for each matrix, their source (SPRINT or other) and remarks (e.g. on coverage of the SPRINT long-list of pesticides). For data generated within SPRINT: all samples were collected during the growth season of 2021.

**Table S1.** Occurrence data sources used in step A.

| No | Matrix | WF[a)] | No. of samples | Source of data | Remarks |
|---|---|---|---|---|---|
| **1a** | Blood serum | 1.0 | 180[b)] | SPRINT (EU) [c,d)] | LC-multi & GC-multi but not full SPRINT scope |
| **1b** | Blood plasma | 1.0 | 140 | SPRINT (EU) [c)] | only LC-multi; only NL and PT |
| **2** | Urine | 1.0 | 2000 | HBM4EU | SPECIMEn study, sampling 2020, NL, CZ, LV, ES, HU [g)] |
| **3** | Feces | 1.0 | 629 | SPRINT (EU) [c)] | does not yet include glyphosate/AMPA |
| **4** | Food | 1.0 | 88,000 | EFSA, 2022a | Sampling in EU (2020), mostly raw agricultural products |
| **5** | Indoor dust | 0.5 | 167 | SPRINT (EU) [c)] | n/a |
| **6** | Air (PUF/PEF) | 0.5 | 20 | SPRINT (EU) [c,e)] | LC-multi & GC-multi but not full SPRINT scope |
| **7** | Soil | 0.25 | 201 | SPRINT (EU) [c)] | n/a |
| **8** | Surface water | 0.25 | 58 | SPRINT (EU) [c)] | n/a |
| **9** | Sediment | 0.25 | 97 | SPRINT (EU) [c)] | n/a |
| **10** | Crops [f)] | 0.25 | 195 | SPRINT (EU) [c)] | n/a |

a) matrix-weighting factor used for overall score, see also section on overall score below; b) samples were pooled from 3-6 subjects of same subpopulation; c) Sample collection SPRINT CSS was done in 2021, sample analysis in 2021/2022; d) analysis outsourced (Limoges, FR); e) analysis external (Kwalis, DE); f) crops from CSS fields, at time of harvest, edible part; g) Huber et al., 2022; Ottenbros et al., 2022.

In the exposure/occurrence assessment, emphasis was on frequency of findings of specific pesticide residues in the various compartments. Concentrations (if readily available) were considered to some extent but not exhaustively. This was done to keep the efforts manageable, to avoid complications due to method LODs/LOQs and dealing with left-censored data (data below a LOD for which the true value is unknown), and also because at this stage the effect of concentrations in the various compartments for human health is not yet clear. The data on frequency of detection and, if possible/available, concentration indications, were compiled in an excel document. Based on these data, for each pesticide an overall score was determined using summation and weighting factors as described below. The higher the overall score, the higher the presumed relevance regarding human exposure.

**Overall score: rough indication of overall (potential) contribution to exposure**

Humans might be exposed through several routes. Measurement of pesticide and their metabolites in human matrices (blood, urine, feces) provides information on aggregated exposure irrespective of the route of exposure. This is therefore considered a very strong indicator for actual exposure. However, since many pesticides are rapidly metabolised and excreted, it typically only reflects recent exposure (1h-24h, depending on the matrix and toxicokinetics of the pesticide). In addition, especially in urine, most pesticides are excreted as (multiple) metabolites. These are not always known and/or analytical reference standards are lacking, and therefore cannot all be covered in the analysis. For this reason, also external exposure data were included in the overall score. Here food is the most important indicator as e.g. fruit, vegetables and cereals may contain relatively high amounts of pesticide (0.01-1 mg/kg), and the oral intake is considerable. Other routes of exposure are inhalation of pesticides in air (pesticides in gas phase or adsorbed to fine dust particles). For these routes, measurements in air and indoor dust were considered indicative. Finally, dermal uptake may occur through skin contact with all sorts of surfaces contaminated with pesticides. The inhalation and dermal routes might be more relevant in terms of relative contribution for farmers and potentially their neighbours.

As far as exposure and occurrence were concerned, the prioritization / pesticide selection was based on the entire population, i.e. no separate selections are being made for individual CSSs, and also not for consumer, neighbour and farmers, and for organic and conventional subgroups.

For each pesticide from the SPRINT long-list, an overall score was determined. In the overall score all routes were taken into account, but differently weighted. HBM was given a relatively high weight, by summation of the frequencies of each of the four measurements (1a, 1b, 2, and 3, see Table 1). To this, frequencies in food were added as such. The pesticide frequencies of the two inhalation-related matrices were not summed as such, but each weighted using a factor of 0.5, to prevent that these would have a higher relative contribution than food. The pesticide frequencies of the four other environmental matrices were each weighted using a factor of 0.25, again to prevent a relative contribution higher than food.

This way, in case of equivalent detection frequencies, the relative contribution for each matrix/route type corresponds to: 4 : 1 : 1 : 1, for [HBM] : [food] : [air(proxies)] : [environment other], respectively. Where information on concentrations (ranges, medians) was available, this information was taken into account and weighting factors for concentration (wfc) were used: for very low concentrations, typically <LOQ, the frequency was multiplied by a wfc of 0.1. For intermediate concentrations a wfc of 0.33 was used, for the higher concentrations no wfc was used (wfc=1). Wfc's were used in all cases except for pooled blood serum samples (concentrations might be reduced due to dilution), and the HBM4EU urine data and the EFSA food data as no concentrations were available in these cases. The overall score was calculated as follows:

Overall SCORE = score [HBM] + score [food] + score [indoor dust & air] + score [soil/water/sediment/field crop

with: SCORE [HBM] = [1a] + [1b*wfc(1b)] + [2] + [3*wfc(3)][JH18.1]

[1a] = detection frequency of PESTICIDE in pooled blood (serum) (%) (SPRINT)

[1b*wfc(1b)]. 1b = detection frequency in blood (plasma) (%); wfc(1b) = weighting factor for concentration: 0.1, 0.33 and 1 for concentrations <0.01, 0.01-0.125 and ≥0.125 ng/mL, respectively.

[2] = detection frequency of PESTICIDE in urine (%) (SPRINT & HBM4EU/SPECIMEn)

[3*wfc(3)]. 3= detection frequency in feces (%); wfc(3) = weighting factor: 0.33 and 1 for concentrations <1 and ≥1 ng/g, L, respectively.

SCORE [food] = [4]

[4] = detection frequency of PESTICIDE in food (%) (EFSA)

SCORE [indoor dust & air proxies] = [WF5*[5]*wfc(5)] + [WF6*[6]*wfc(6)]

[WF5*[5]*wfc(5)]: WF5 = matrix-weighting factor of 0.5; [5] = detection frequency in indoor dust (%); wfc(5) = weighting factor for concentration: 0.1, 0.33 and 1 for concentrations concentrations <50, 50-250 and ≥250 ng/g, respectively. (SPRINT)

[WF6*[6]*wfc(6)] = WF6 = matrix-weighting factor of 0.5; [6] = detection frequency in air proxies (passive sampler, PUF or PET) (%); wfc(6) = weighting factor for ng/sampler: 0.1, 0.33 and 1 for <50, 50-250 and ≥250 ng/sampler, respectively. (SPRINT)

SCORE [soil/water/sediment/crop] = [WF7*[7]*wfc(7)] + [WF8*[8]*wfc(8)] + [WF9*[9]*wfc(9)]+ [WF10*[10]*wfc(10)]

[WF7*[7]*wfc(7)] = WF7 = matrix-weighting factor of 0.25; [7] = detection frequency in soil (%); wfc(7) = weighting factor concentration: 0.1, 0.33 and 1 for concentrations <50, 50-250 and ≥250 ng/g, respectively. (SPRINT)

[WF8*[8]*wfc(8)] = WF8 = matrix-weighting factor of 0.25; [8] = detection frequency in water (%); wfc(8) = weighting factor for concentration: 0.1, 0.33 and 1 for concentrations <0.01, 0.01-0.1 and ≥0.1 µg/L, respectively.

[WF9*[9]*wfc(9)] = WF9 = matrix-weighting factor of 0.25; [9] = detection frequency in sediment (%); wfc(9) = weighting factor for concentration 0.1, 0.33 and 1 for concentrations <1, 1-10 and ≥10 ng/g, respectively.

[WF10*[10]*wfc(10)] = WF10 = matrix-weighting factor of 0.25; [10] = detection frequency in field crop (%); wfc(10) = weighting factor for concentration 0.1, 0.33 and 1 for concentrations <10, 10-50 and ≥50 ng/g, respectively.

Output of step A: ranking of all 206 SPRINT Pesticides based on their occurrence score, to identify the top 25 Pesticides in terms of exposure relevance. This short-list of 25 was used as input for step B, potential hazards for human health.

### Step B: Health hazard

In this step the top-25 list of highest prioritised Pesticides based on occurrence was used. In this step the hazard will be taken into account as well. A hazard of Pesticides is defined as an intrinsic toxic property of a pesticide active ingredient. For this procedure its relevance to humans is key.

Not all hazards are relevant to all humans: e.g. irritancy or sensitization for dermal or inhalation routes may be relevant to farmers and other applicators and perhaps also to some extent for neighbours but certainly not to consumers who are primarily exposed to pesticide through the diet or by occasional home applications of Pesticides and/or biocides. So, farmers and neighbours are seen as consumers with additional exposure related to direct and indirect non-dietary exposure related to multiple routes (via air, via tracking, etc.). Table 2 provides an overview of the use scenarios and proposed hazard evaluation framework for hazard ranking.

Here we will try to identify the most critical adverse effect for each pesticide based on existing data. The critical hazard is defined as 'the lowest no effect level relevant to human health'. This results in a low tolerance level for exposure. For the human health hazard of pesticide the ADI and ARfD could be used for long-term and short-term dietary exposure, respectively. For non-dietary sources and routes of exposure we will to rely on existing procedures for health-based recommended occupational exposure limits (OELs). For this, EFSA introduced the Acceptable Operator Exposure Level (AOEL) for long-term and Acute Acceptable Operator Exposure Level (AAOEL) for short-term exposure.

We will prioritize components that have been classified by IARC as group 1 ("carcinogenic for humans") or group 2A ("probably carcinogenic for humans") and the criteria for pesticides as 'highly hazardous'. Here we will also consider the use of weight factors that will reflect relevance to humans, e.g.:

- Recent systematic evaluations of human evidence, including SPRINT pesticide effects on human, animal and ecosystem – systematic scoping reviews (deliverable 3.1)
- In vivo studies to provide evidence supporting biological plausibility
- Mechanistic/molecular evidence from in vitro studies to provide supportive mechanistic evidence

- For certain adverse outcomes there may be published adverse outcome pathways (AOPs) and key characteristics (KCs) that are considered to inform mode of action in humans (Guyton et al., 2018; La Merrill et al, 2020; Smith et al., 2020)
- EFSA reports (EFSA, 2019a, 2019b, 2022a, 2022b, 2022c)

Exclusion criteria: Hazardous pesticides that are currently banned or severely restricted in EU will be excluded.

Output: hazard assessment of the top 25 Pesticides from step A.

**Table S2.** Hazard ranking for the general population and the worker's populations.

| Population | General population | Occupational population |
|---|---|---|
| **Description** | Consumers, residents and bystanders | Persons who receive additional work-related exposure e.g. during pesticide applications |
| **Exposure scenarios** | Dietary intake and indoor and outdoor use in a residential setting, also including intrusion of pesticides by air, dust or other causes including residential and bystander exposure as defined by EFSA (e.g. EFSA 2014) | All indoor and outdoor application of pesticide products and exposure after application (e.g. dermal exposure after re-entry and other exposure scenarios, e.g. defined in EFSA (2022a) |
| **Hazard ranking** | Existing framework of reference:<br><br>IARC Classification (Group 1 or 2A)<br><br>Acceptable daily intake (ADI)<br><br>Acute Reference Dose (ARfD) | IARC Classification (Group 1 or 2A)<br><br>Acceptable Operator Exposure Level (AOEL) and Acute Acceptable Operator Exposure Level (AAOEL) or existing TLV for 8-h or 15-min in one or more EU countries (e.g. listed at https://limitvalue.ifa.dguv.de/ |
| | Only if existing guidance values are not available such values will be derived from NO(A)ELs from animal studies, preferably a 90-d | Only if existing AOELs are not available we will use NO(A)ELs from animal studies to set provisional AOELs (EC, 2006), taking into account short-term and/or long-term (working life) |

| | | |
|---|---|---|
| | study (ECHA, 2016). For the selection of the critical effect, we will take into account population-based studies identifying certain adverse health outcomes in vulnerable subgroups (e.g. effects during pregnancy, effects in early childhood or at old age). For this the information provided in D3.1 can be used. | exposure duration. The critical effect will be selected based on the relevant use scenario (to determine the main route of uptake). For prioritization human population-based studies identifying certain adverse health outcomes will be considered. For this the summary from a recent systematic scoping of the literature will be used. |

**Pesticide hazards for humans**

The hazard of a substance (or mixture) is defined as the intrinsic property to cause adverse effects. For the pesticide selection we preferably use human data. If these are not available or cannot be used, data from toxicity studies in animals (so-called in vivo studies) are also available. It is important to verify to what extent the animal model is a good model for predictions of toxic hazard in humans.  For the current analysis we considered which of the observed adverse effects is the most 'critical effect' in a given exposure scenario for different groups in the population. The critical effect is the first effect observed at increasing dose for which there is consensus that this effect may be considered adverse to health. The occurrence of this critical effect may depend on the (primary) route of exposure and the exposure level. Below we will discuss the main exposure scenarios of consumers (C) and of farmers (F) and their families and other inhabitants of the rural environment who may live close to agricultural fields where Pesticides are applied. The last group is referred to as 'neighbours' (N). Farmers and neighbours (F+N) are discussed together because their exposure scenarios are similar in mechanisms (although their degree of exposure may differ substantially). In the terminology used by EFSA the neighbour is covered in the 'bystander' definition (EFSA, 2022b). Family members of the farmer household are also considered bystanders (ref). Below the exposure scenarios of consumers, farmers and neighbours will be discussed in more detail.

Consumers (C) will be primarily exposed to Pesticides from the diet and to some extent also from home and garden use of a specific Pesticides as e.g. biocide or veterinary use on companion animals. The exposure of C depends on the absorption of pesticide residues from food in the gut. There may also be other types of contact, e.g. food preparation. Indirect exposure is to be expected from contamination of the indoor environment due to intrusion of outdoor Pesticides in the gas phase or airborne (fine) dust. Homes may also become contaminated from clothing and footwear (so-called tracking). Exposure can be explained by direct skin absorption or indirectly by so-called hand-mouth behaviour (known as finger shunt) that can be observed in (young) children. The fraction that is absorbed from the gut will be transported to the liver by the hepatic portal vein (HPV) where it will be (further) metabolized. For most pesticide active ingredients, the liver metabolism will lead to products that are less toxic than the parent. The safety of C is protected by maximum residue limit (MRLs) on pesticide residue levels in food items. To cover the intake of pesticide residues from different food items in the daily menu there are two additional reference values: for short-term effects the ARfD is used and for long-term exposure the ADI is used as a reference for as the amount taken up that is considered safe over a lifetime exposure. If this value is low. This means that the toxicity for C is considered to be high and vice versa.

Farmers (F) may be exposed to Pesticides by food and direct/indirect routes in their homes (see above), and additionally from direct contact with Pesticides during their involvement in pesticide applications or exposure to the residues on the crops upon re-entry (when working in the sprayed field). This occupational exposure is studied for many Pesticides and is known to be an important source of uptake. For the assessment of risk, it is important that exposure routes may be different compared to C. When applying Pesticides, inhalation and skin may be significant additional routes of uptake that may explain a large part of total uptake. Inhalation of Pesticides in the gas phase as a liquid aerosol or associated to dust particles may lead to uptake through the airways. This is a fast process and a concern if Pesticides contain active ingredients and co-formulants harmful to the airways. Inhalation also leads to fast and systemic uptake. In addition, farmers may be dermally exposed as a result of direct contact of the unprotected skin. Dermal absorption is a much slower process (compared to inhalation) and health effects and may occur with a delay. There are many pesticide active ingredients that contribute to the body burden by skin uptake: when wearing respiratory protection this may become a major route of uptake. Indirect exposures may also occur by ingestion, e.g. following hand-to-mouth contact as a result of poor personal hygiene practice but this is not a main source of exposure when farmers are well aware of the risks and protect themselves. To protect workers from adverse health effects occupational exposure limits (OELs) are used. After Pesticides are taken up via inhalation and skin, they can reach all internal organs depending on their ability to cross the blood brain and the blood placenta barriers. To some extent Pesticides can be detoxified by different tissues but the most important contribution is from liver metabolism. For this Pesticides have to be delivered to the liver. Reference levels for inhalation such as occupational exposure levels (OELs) are based on the most critical health hazard related to the specific exposure scenarios that are anticipated when applying Pesticides that are more likely to be related to inhalation and effects on the airways. There are many Pesticides that are more harmful when inhaled compared to uptake by ingestion or following dermal absorption. Therefore, the overall risk assessment for farmers is different, and the pesticide selection for farmer's exposure and can deviate from the selection for C. Neighbours (N) are not using Pesticides professionally. However, due to their residential setting which may be close to agricultural land where Pesticides are applied, they may become exposed during spraying by the drift of liquid aerosols and vapor or later due to evaporation of Pesticides from crops. In addition, there are secondary routes of exposure to Pesticides such as from wind-blown soil dust, surface water or taking home of dirt under shoes (tracking). Neighbours may thus have exposure by additional sources and routes of exposure compared to the 'average' consumer. This exposure is much lower than occupational exposures. However, the health risks are different as explained above, also because the route of uptake may be different from that of the consumer. Most of the oral exposure is probably from dietary intake and therefore the ADI and ARfD can be used as a framework of health risk assessment. Due to non-dietary exposure, it is suggested to use the framework of health risk assessment of the farmer and also the dose metrics considered to protect the farmer. A limitation of this use of OELs is that they assume exposed people to be of working age and do not consider susceptible people, e.g. young children or elderly and other people who have higher susceptibility such as during pregnancy. In the risk assessment practice this higher vulnerability is often covered by an assessment factor of 10 to cover interindividual differences in susceptibility within the human population. This means that the OEL should be divided by a factor of 10 or more, depending on what is known about potential increased susceptibility.

**Table S3.** How biomonitoring data can inform adverse health effects of pesticide exposure.

| Biological matrix | Target organ | Toxicity mechanism | Adverse health effects attributed to pesticide |
|---|---|---|---|
| **Blood** | Central and peripheral nervous system | Neuro apoptosis, interactions with receptors of neurotransmitters, and other neurotoxic mechanisms | Neurodevelopmental and neurodegenerative disease (Parkinson's disease, ADHD, Autism, mental health) |
| | Bone marrow | Genetic and epigenetic changes to chromosomes | Non-Hodgkin lymphoma and leukemia in adults and children |
| | Fetal exposure following placenta passage | Neurotoxicity, oxidative stress, [other] | Anomalies at birth with potential consequences later in life |
| | Endocrine system | (Weak) interaction with hormone receptors | Sub/infertility |
| **Urine** | Kidneys | Oxidative stress, nephrotoxicity, [other] | Chronic kidney disease |
| **Feces** | Gastro intestinal tract | Changes in microbiome and their potential health effects (e.g. gut-brain axis hypothesis) | Neurodevelopmental and neurodegenerative disease |
| **Nose mucous** | Upper airways | Inflammatory response, oxidative stress, irritative and allergic responses (elicitation, sensibilization) | Airway infections, airway response, lung function changes, asthma |

Linking exposure to health effects

Table S3 provides an overview of how concentrations in biological media can inform target dose and which toxicity mechanisms are involved. To be sure that we cover all human health endpoints of interest for the SPRINT project a flow diagram was prepared how the occurrence data from Step A are used in Step B to calculate the hazard quotients (Figure 2). In this flow chart we start with identifying the Pesticides that have only a classification for toxicity related to a local effect e.g. on skin or eyes (H310-319) or on airways (H320-335). If this is the case, Pesticides should be considered for testing in a suitable in vitro test, i.e. a test system for effect on eyes, skin and/or airways. For these Pesticides inhalation exposure data are required that could be derived from pesticide levels in gas phase or airborne particles e.g. from air sampling close to the site of application. In this subgroup priorities for pesticide selection may be set based in concentration levels in air (either gas or particle phase or both).

If the substances do not have a local effect or have other effects next to a local effect this takes us to the section in the flow diagram that describes systemic effects. Here we start with the simple scenario that a pesticide active ingredient that is taken up by ingestion passes through the GI tract and reaches the gut where it may have an effect on the gut microbiome. Note that if substances are found in feces this could also be the result of excretion from the body by bile or pancreatic excretion, but this cannot be concluded without considering the results of analysis of blood and/or urine.

If pesticide active ingredients are not reported in feces the next step is to find out if they were detected in blood fractions. If yes, the substance qualifies for in vitro and in vivo testing related to potential toxicities affecting one or more internal organs or organ systems. Due to short half-life in blood, it is possible that Pesticides are not detected in a single random (snapshot) blood sample. Therefore, we also consider the occurrence in urine.  Here the pesticide may be found as a parent but much more likely as a metabolite. For urinary excretion of metabolites to indicate exposure to a pesticide the metabolite should be specific for a pesticide, or if not specific additional information such as a detect in plasma or feces or external exposure sources supports a positive lab result based on urine analysis. All substances that are entering Step B will be part of the list of Pesticides ranked by HQ.

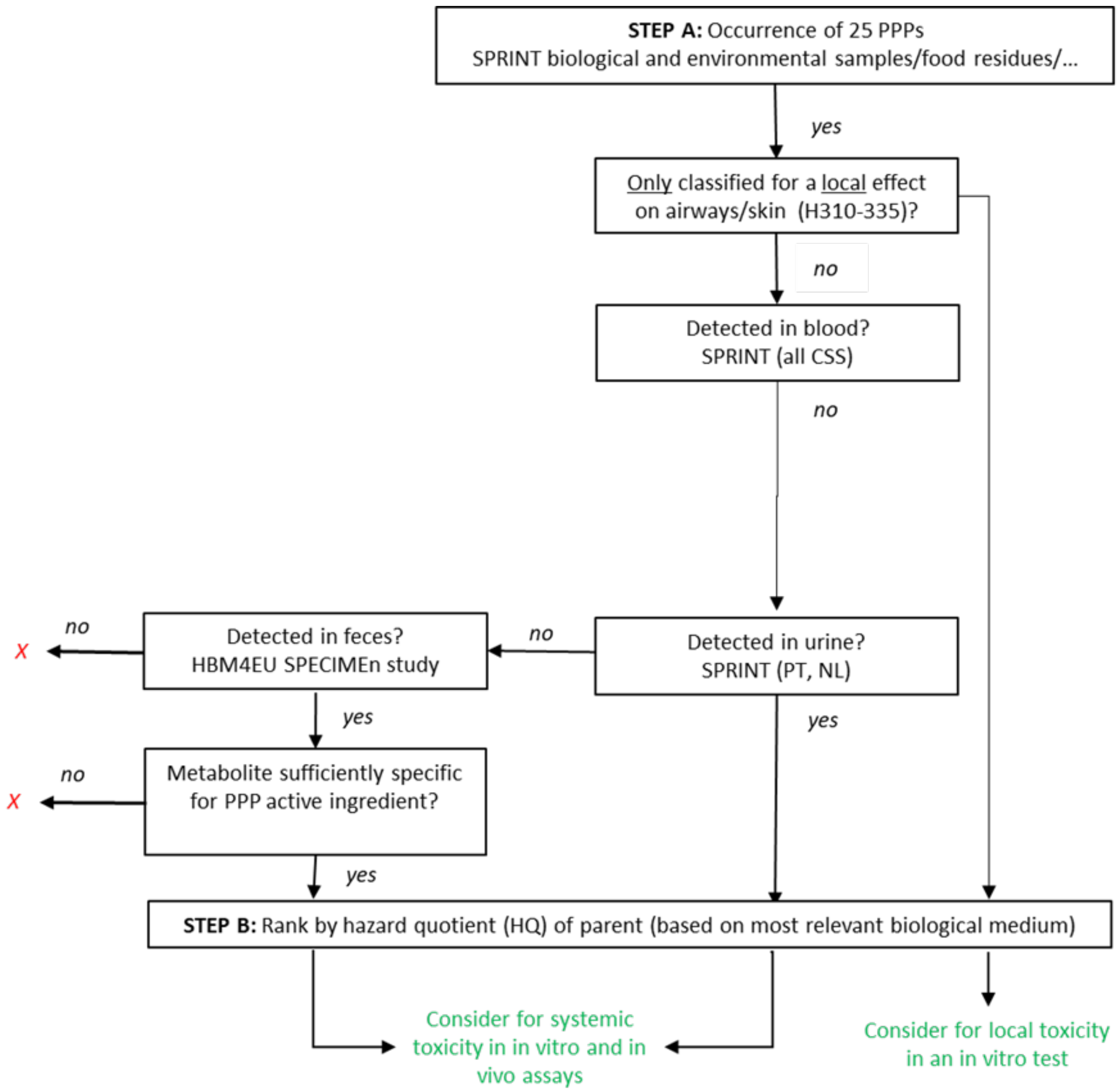


**Figure S2.** Flow diagram showing the process starting from the short list of 10 pesticides (Step A) and how available hazard classifications and biomonitoring data are used to perform to complete the ranking process based on hazard (Step B).

**Calculations of the weighted hazard quotient**

Hazard quotient (HQ) is the ratio of the potential exposure to a substance and the level at which no adverse effects are expected.

For **farmers**: HQ = PESTICIDEL / AOEL (also used for **neighbours**)

For **consumers**: HQ = PESTICIDEL / ADI

With: PESTICIDEL = PESTICIDE level (mg/kg bw/day)

AOEL = Acute Occupational Exposure Level (mg/kg bw/day)

ADI = Acceptable Daily Intake (from food) (mg/kg bw/day)

w = weighting factor by detection rate

**Conversions**

Inhalation - Average inhalation volume for 8 hours is 10 m3 (light exercise). A workshit of 8 is converted to 24 hours by multiplying with a factor of 3. For an average person's body weight for adult (M/F) we used 70 kg. This leads to the following conversion:

OEL (mg/m3) x 10 (m3) x 3 / 70 kg bw => mg/kg bw/d

Blood - Conversion of serum and plasma concentrations in µg/L to mg/kg bw/d (for ADI and AOEL). In this conversion the average blood volume in an adult (M/F) was assumed to be 6.0 L and an average body weight for an adult (M/F) is 70 kg

Plasma (µg/L) x 6 L / 70 kg bw / 1000 => mg/kg bw/d

Additionally, for the pooled serum sample (analysed in Limoges): one sample is diluted by two others = correction factor 3 x (worst case):

Serum content (µg/L) x 3 x 6 L / 70 kg bw / 1000 => mg/kg/d

Feces - Conversion of feces concentration in µg/kg feces to mg/kg body weight/day (mg/kg bw/d) for ADI and AOEL. Assuming an average production of feces per day of 0.5 kg and an average body weight for adults (M/F) of 70 kg this gives the following conversion equation:

Feces content (µg/kg) / 70 /2 / 1000 => mg/kg bw/d

Note: these conversions are crude and can only be used for prioritization (not for quantitative risk assessment purposes). The plan is to refine conversions by PBK modelling (in WP3) and make sure that we know how accurate these conversions are and how they may be dependent on pesticide and host-factors (e.g. biotransformation, differences by gender, etc.)

#### Step C: Synthesis

The top 10 Pesticides from step A will be plotted in a risk matrix based on an index in arbitrary units (to reflect occurrence and exposure on X and the lowest 'critical' hazard expressed as mg/kg body weight on the Y-axis. In this graphic presentation by consensus, we will identify potential pesticide mixtures for toxicity testing based on co-occurrence, co-exposure and inferences regarding mode of action and reported observed interactions (EFSA 2019a, 2019b). To generate a risk matrix plots we will use an online tool https://risk21.org/webtool/. The results are shown in **Figure S3** and **Figure S4**.

We realize that the standard solution offered uses an 'estimate of exposure'. In our case we will use an integrated expression combining different types of information on occurrence and exposure with a weight factor that will be expressed as an index in arbitrary units and lead to ranking the pesticides.

The short list of 10 pesticides included: glyphosate, acetamiprid, tebacunazole, lambda-cyhalothrin, cypermethrin, deltamethrin, cyprodinil, fluopyram, imazalil, and the synergist piperonyl butoxide (synergist), Despite the differences between HQs based on ADIs (derived from consumer exposures) and those derived from OELs (from farmer's and neighbour's exposures) good agreement was observed in the rank numbers (see **Table S4).**

**Table S4.** Selection of Top-3 and Top-10 for toxicity testing.

| Rank | Pesticides |
|---|---|
| **1** | Glyphosate |
| **2** | Acetamiprid |
| **3** | Tebuconazole |
| 4 | Lambda-Cyhalothrin |
| 5 | Cypermethrin |
| 6 | Deltamethrin |
| **7** | Cyprodinil |
| **8** | Piperonyl butoxide (synergist) |
| **9** | Fluopyram |
| **10** | Imazalil |

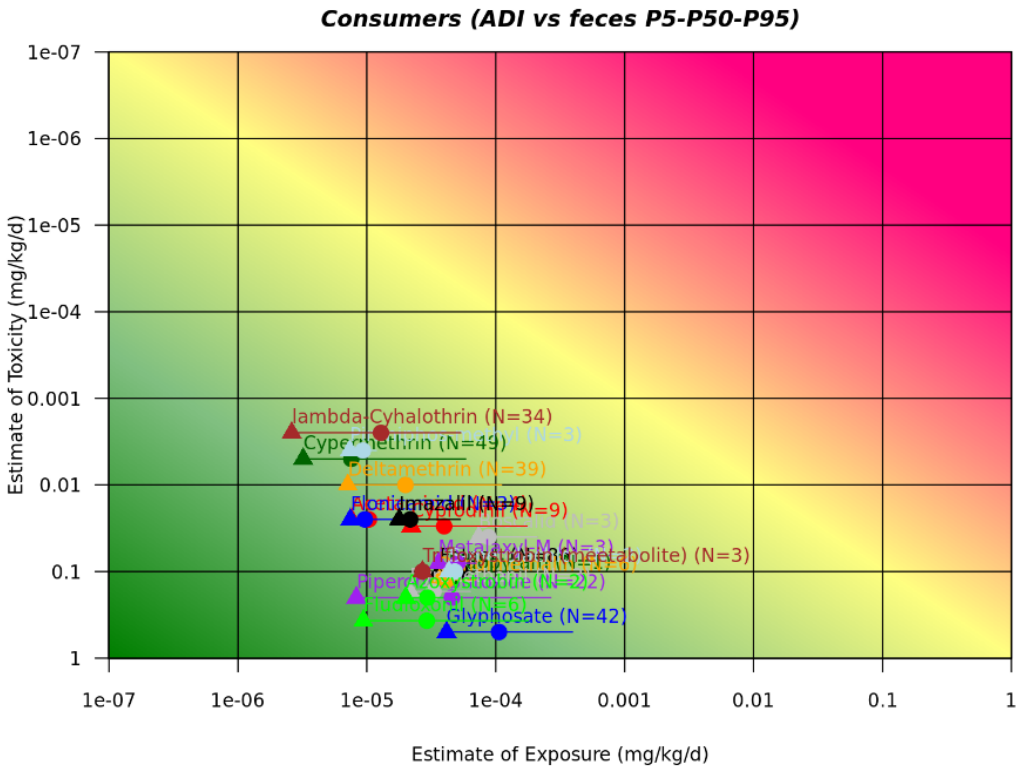


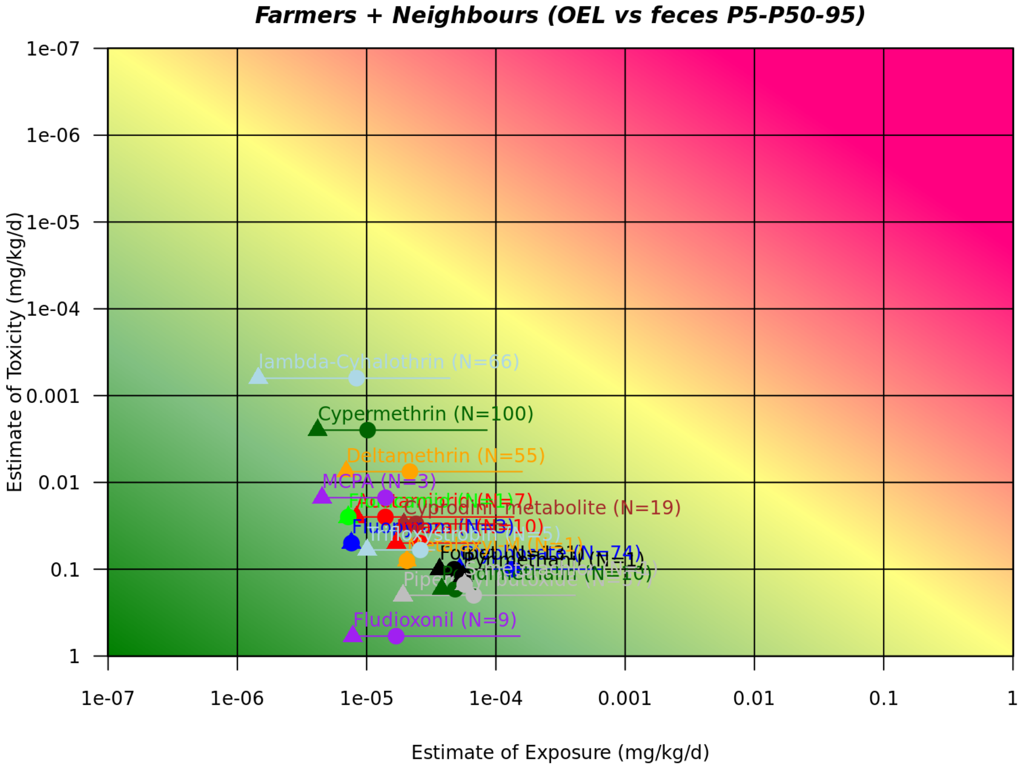


**Figure S3.** Risk matrix based on detects >LOQ in feces of 233 consumers (upper panel) and 396 farmers + neighbours (lower panel) from all CSS. Numbers of detects >LOD are given in parenthesis behind the pesticide names. The total number of different pesticide residues was 12 for consumers and 19 for farmers and neighbours.

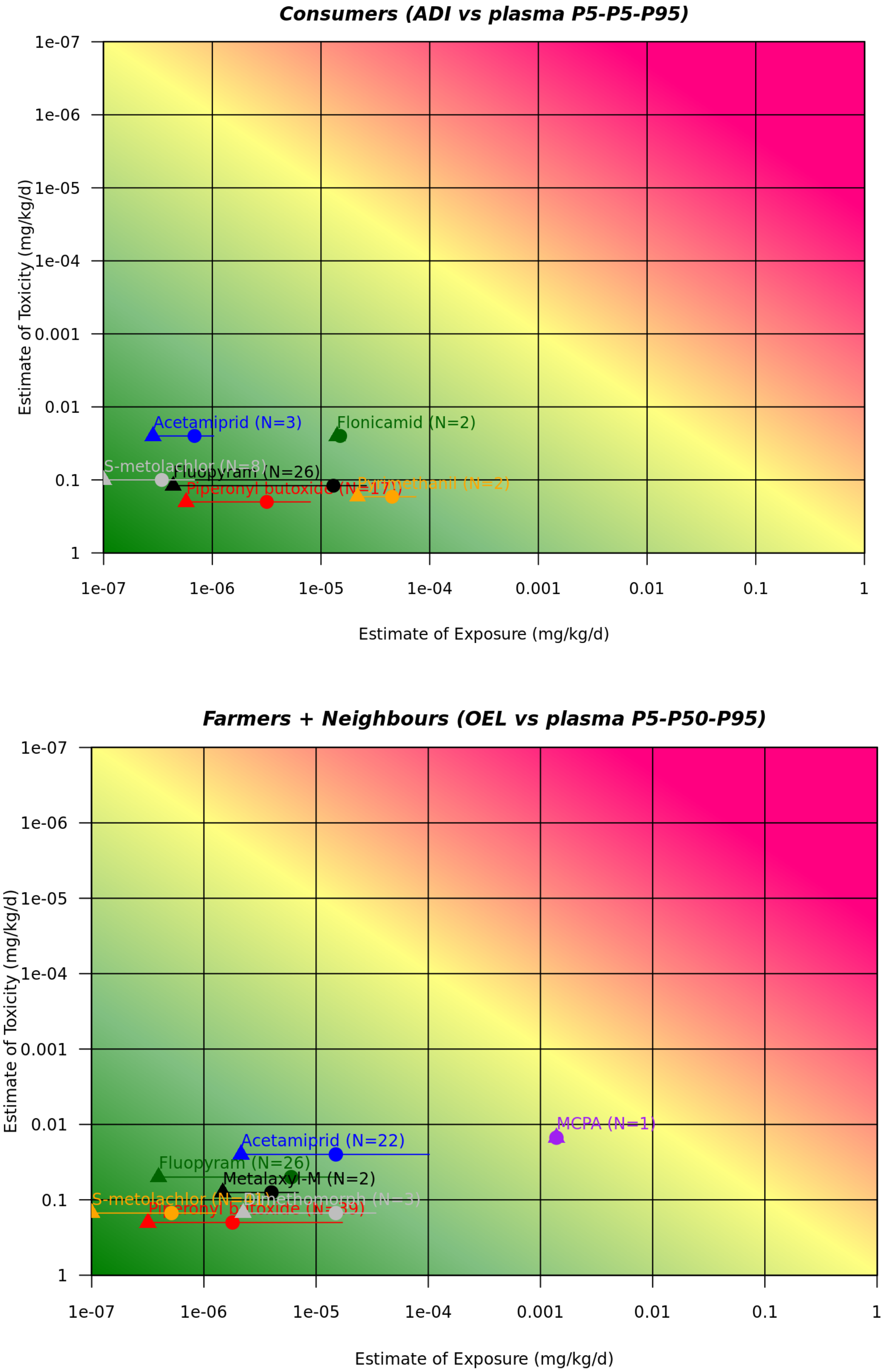


**Figure S4.** Risk matrix based on detects >LOQ in plasma of 47 consumers (upper panel) and 91 farmers and neighbours (lower panel) from all CSS. Numbers of detects >LOD are given in parenthesis behind the pesticide names. The total number of different pesticide residues was 6 for consumers and 7 for farmers and neighbours.
